# Space- and object-based attention in patients with a single hemisphere following childhood resection

**DOI:** 10.1101/2024.12.06.627251

**Authors:** Sophia Robert, Michael C. Granovetter, Shouyu Ling, Marlene Behrmann

## Abstract

The neural processes underlying attentional processing are typically lateralized in adults, with spatial attention associated with the right hemisphere (RH) and object-based attention with the left hemisphere (LH). Using a modified two-rectangle attention paradigm, we compared the lateralization profiles of individuals with childhood hemispherectomy (either LH or RH) and age-matched, typically developing controls. Although patients exhibited slower reaction times (RTs) compared to controls, both groups benefited from valid attentional cueing. However, patients experienced significantly higher costs for invalid trials—reflected by larger RT differences between validly and invalidly cued targets. This was true for invalid trials on both cued and uncued objects, probes of object- and space-based attentional processes, respectively. Notably, controls showed no significant RT cost differences between invalidly cued locations on cued versus uncued objects. By contrast, patients exhibited greater RT costs for targets on uncued versus cued objects, suggesting greater difficulty shifting attention across objects. We explore potential explanations for this group difference and the lack of difference between patients with LH or RH resection. These findings enhance our understanding of spatial and object-based attention in typical development and reveal how significant neural injury affects the development of attentional systems in the LH and RH.

## Introduction

Visual information in the environment is highly complex and unfolds quickly. Attention is a cognitive mechanism that biases an observer to respond rapidly and effortlessly to behaviorally relevant or salient subsets of this visual input^1,2^. This selection of information, which favors the processing of certain locations or objects^3^ at the expense of others^4^, can occur covertly in the absence of overt eye movements. Covert attention is considered to be the output of competitive interactions between bottom-up/stimulus-driven information from the external environment and top-down signals/internal goals of the observer. ‘Spatial-based’ attention refers to the selection and preferential processing of specific *positions* in space, whereas ‘object-based’ attention refers to the selection and preferential processing of specific *objects* (and perhaps also their associated spatial location) over others. Whether these forms of attention arise from the same underlying neural substrate or are, at least to some extent, independent, is not yet fully resolved^1,2^.

### Spatial-based attention is right-lateralized in adults

Spatial-based attention has been well characterized using the covert cueing attention paradigm designed by Posner and colleagues^5,6^. In this paradigm, two squares are presented on a screen, one on either side of central fixation (and participants maintain central fixation). In the exogenous version of the task, a cue – e.g., a brightening of one square – draws attention to that square. If the subsequent target (e.g., an asterisk) appears in the cued square, target detection is facilitated (‘valid’ trials) compared to when the target appears in the uncued box (‘invalid’ trials). The rapid responses to valid trials reflect the benefit of the target being presented in the attentionally-cued spatial location or, in invalid trials, the cost of switching attention to the new location at which the target appears. Valid trials are the most common (up to 70-75%) and this induces participants to attend preferentially to and make use of the spatial location of the cue.

Studies typically localize the neural basis of spatial attention and selection to the right hemisphere (RH) in humans but not in other non-human primates^7,8^. This hemispheric asymmetry in humans is thought to be a consequence of the fact that, in the large majority of the population, the left hemisphere (LH) is dominant for language and, hence, attention is largely mediated by the RH^9,10^. The RH superiority for spatial-based attention has been confirmed in both positron emission tomography^11^ and functional magnetic resonance imaging (fMRI)^12^ studies using paradigms that closely resemble the Posner covert spatial cueing experiment above^13^. Further evidence for this hemispheric lateralization comes from studies of adults with damage to the right parietal lobe^14^, with or without concurrent hemispatial neglect^15^, a condition marked by reduced awareness of the contralesional side of space^16^. Those with RH damage are deficient in spatial-based attentional processing to a greater degree than those with damage to the left parietal lobe^16–19^.

### Object-based attention is left-lateralized in adults

In addition to attending to and selecting particular spatial locations, covert attention can also be directed to particular objects. In the widely replicated two-rectangle paradigm shown in Figure 1^15^, a target that appeared at a location (top or bottom of one of two displayed rectangles) that is validly precued (75% of trials) was detected faster than a target that appeared at an invalid, uncued location (25% of trials)^15^. Critically, however, the reaction time (RT) cost on invalid trials differed depending on where the invalid target appeared (compare rows 2 and 3 in Figure 1). Relative to valid trials, RT was slowed by a further 47 ms when the target appeared in an uncued spatial location on the uncued object (invalid-space condition, IS), demonstrating the cost of spatial-based attention, i.e., shifting attention away from the cued location. However, RT was slowed by only an additional 34 ms when the target appeared in an uncued spatial location but *within* the cued object (invalid-object condition, IO). Note that the IS and IO targets were equidistant from the cue. If only spatial-based attention played a role, RT should have been equal between IS and IO trials. The 13 ms statistically significant advantage when the target fell within the cued (IO) vs uncued (IS) rectangle thus reflects object-based attentional modulation, i.e., the positive contribution that accrues from the shared representation of the cued rectangle. This may reflect an ‘object-based advantage’ of attention spreading within the cued object, resulting in faster RTs on IO trials^20,21^, and/or an additional ‘object switching cost’ that arises when attention shifts across different objects, leading to slower RTs on IS trials.

**Figure 1.**
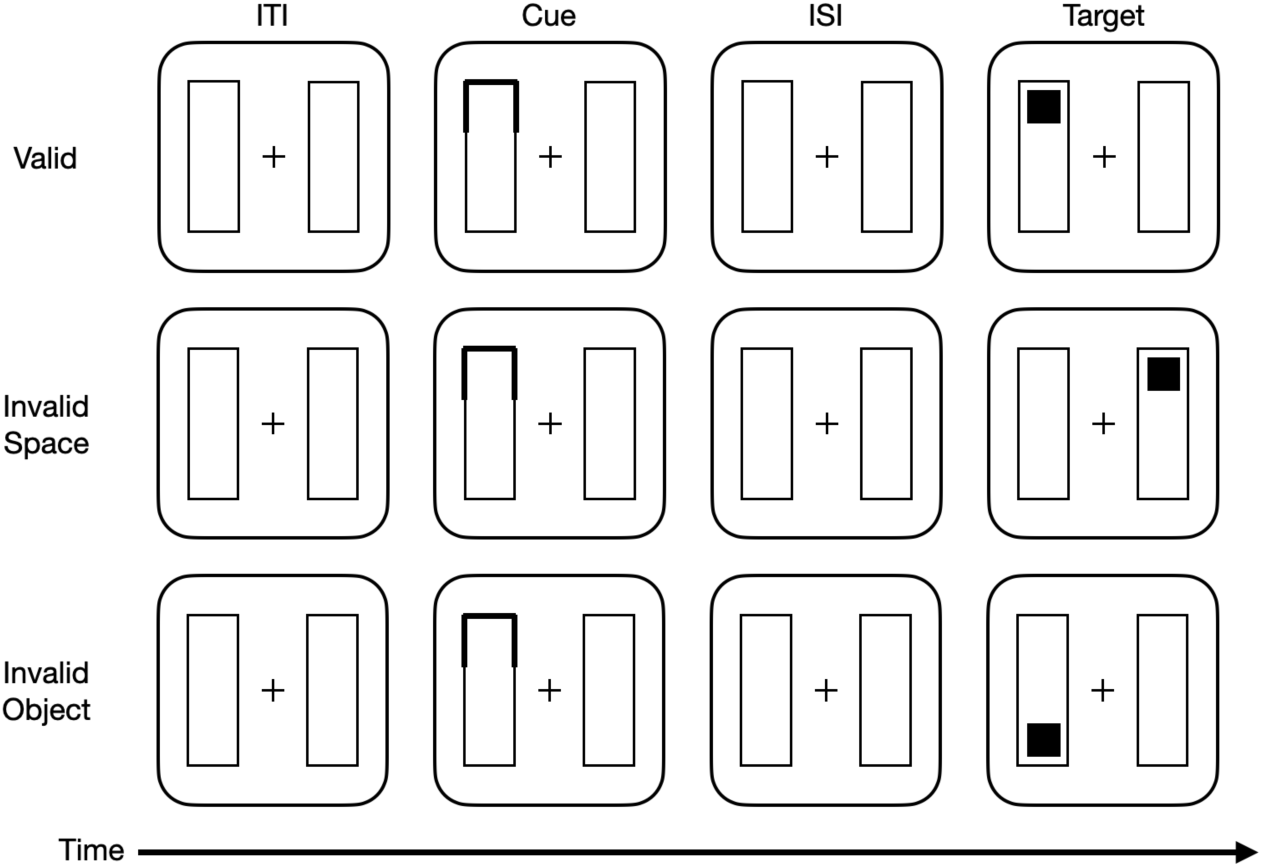
The two-rectangle attention paradigm of Egly, Driver, et al., 1994. Two rectangles appear on the screen, followed by an ‘exogenous’ or bottom-up cue (black outline), in one of four locations. After a brief pause, usually 100-150 ms, the target (black square) appears at one end of one rectangle in one of three possible locations (never occurs on the diagonal opposite the cued corner). In *Valid* (V) trials, the target appears in the same location and object (rectangle) as the cue – this occurs on roughly 70% of the trials, leading the observer to use the cue to predict target location. In the *Invalid Spatial* (IS) trials (second column), following the cue, the subsequent target appears at a different location, at the same relative position on the other, uncued object. In the *Invalid Object* (IO) trials (third column), following the cue, the subsequent target appears at an uncued location that is spatially equidistant from the cue as in IS trials but is within the same cued object. *Neutral* (N) trials (fourth column), are the baseline condition in which all four ends of the rectangles are cued, offering no prediction of upcoming location of the target.

In a seminal study implicating the LH in object-based attention, Egly, Driver, and colleagues^15^ used this paradigm to localize space- and object-based attention in a split-brain patient, in whom communication between the hemispheres is not possible due to resection of the corpus callosum. By presenting stimuli to one hemifield at a time, and ensuring that the patient maintained central fixation, they were able to evaluate the performance of each hemisphere independently. This patient displayed significant differences in RT costs for IS and IO trials, where costs are defined as the RT on invalid trials relative to those on valid trials, when the display was in the left hemifield (RH). By contrast, when the display was in the right hemifield (LH), there was a significantly greater cost for IS trials than for IO trials. This finding suggests that the LH, operating in isolation, has an additional cost when the target appears in the uncued object, requiring a switch of attention across objects. As this sensitivity to cued versus uncued objects was not observed for the RH, these results highlight a greater role of the LH than for the RH for object-based attention and its modulation of spatial costs.

Further support for the LH involvement in object-based attention comes from a neuroimaging study. Shomstein and Behrmann^22^ employed a modified version of the Egly paradigm in a group of healthy adults, requiring them to perform object- and space-based shifts of attention while undergoing fMRI. As anticipated, right posterior parietal cortex activity was associated with mediating spatial shifts of attention for both IS and IO trials, since the distance from the cue was equal in these conditions. More notably, only the left posterior parietal cortex exhibited greater activation for shifts to IO targets compared to IS targets, indicating a lateralization of object-based attention to the LH in adults.

### Spatial- and object-based attentional processing in childhood

While significant progress has been made in documenting the behavioral differences between space- and object-based attention and their hemispheric lateralization in adults, much less attention has been paid to whether such processes are already present in childhood and how they are neurally organized. In the studies that do exist, different operational definitions and experimental assays of “attention” have made it difficult to reach a consensus about the neural correlates of spatial and object attention in children. For spatial attention, a mix of right-lateralized and bilateral activation patterns have been observed depending on task and age. Some studies have reported bilateral processing of spatial information during visual-spatial construction and reconstruction across a range of ages^e.g., 23,24^. In contrast, others have found right-lateralization during tasks involving spatially complex visual search, with better performance correlating with higher degrees of lateralization, a relationship that increased with age^25,26^. In studies that relied on paradigms more closely related to the study conducted here, i.e., in which valid and invalid targets are compared, covert shifts of attention have been reported in 4-month-old infants^27^, and rapid development of spatial attention has been observed from 5-10 months of age^28^. No object-based modulation of spatial effects was investigated in these studies.

Some useful evidence is also offered by neuropsychological studies of attention (again, largely space-based) in children with isolated RH or LH damage, although these studies have also produced mixed results. There are reports of deficits in children with cerebrovascular injury in engaging or orienting attention^29,30^, as well as subtle, persistent attention biases in children with perinatal injury, regardless of which hemisphere is affected^31^. Hemispatial neglect can also occur in children with brain damage; however, its manifestation in children is highly variable and not consistently linked to RH versus LH damage^32^. For instance, in a cohort of 34 pediatric hemispheric surgery patients, only one was reported to have neglect^33^.

In a large study, Adamos and colleagues^34^ showed that children with LH perinatal stroke showed extensive impairments in orienting and disengaging attention to both sides of visual space, while those with RH perinatal stroke had more limited impairments, confined to the contralateral side. This is the opposite of what one may predict on the basis of the adult profile RH specialization for spatial attention^35^. On the other hand, another study has shown that there are hemispheric differences in attentional capacities in children with unilateral brain damage. Danguecan and Smith^36^ conducted a presurgical evaluation of 91 children with LH epilepsy that showed that the necessitation for language function to be accommodated in the RH can compromise ‘native’ functions of the RH such as attention. Patients with RH language dominance had poorer scores on visuo-spatial measures compared with patients with more typical LH dominance. Despite these findings, none of the studies clearly differentiate between spatial- and object-based attention, leaving open the question of when hemispheric specialization for these attentional processes begins to emerge and what consequences ensue if one hemisphere is resected in childhood.

### The current study

Here, we investigate how development with only one hemisphere (left or right) affects spatial and object-based attention. One reason that hemisphere-specific deficits might be observed less commonly in children with cortical damage ^32^ compared to adults might be because children, in general, have relatively less lateralized spatial- and object-based processing. Hence, damage to a single hemisphere may be compensated for by intact attentional functions in the opposite, undamaged hemisphere. Another reason is that, given the enhanced plasticity during childhood ^35,37,38^, children may be able to compensate for deficits in lateralized functions via plastic processes regardless of which hemisphere was initially affected. Alternatively, it is possible that when the entire cerebral hemisphere is removed, hemisphere-specific pressures will differentially affect the development of spatial- and object-based processing. To test these hypotheses, we recruited individuals who underwent the surgical removal of an entire hemisphere (hemispheric surgeries, including hemispherectomy and hemispherotomy,^39^ of the LH or RH) during childhood.

First, we compared the patients’ spatial- and object-based attention against that of age-matched typically developing (TD) controls using a child-appropriate Egly two-rectangle paradigm. Second, we compared patients with only a LH to those with only a RH to test whether the hemisphere resected differentially affects spatial- vs. object-based attention. The answer to these questions will be informative with respect to our understanding of typical development of attention, the potential for cortical plasticity and, perhaps, even for the enhancement of attentional functions.

## Methods

### Participants

21 left (8 female, age range: 5.53 – 30.08 years old) and 14 right (8 female, age range: 5.06 – 32.41 yr old) childhood hemispheric surgery patients and 24 TD age-matched controls (14 female, age range: 6.49 – 32.41 yr old) were tested. We provide detailed demographic information about patients in Table S1. Because of the persistent hemianopia^39^, patients viewed the paradigm entirely in their intact visual field. To match presentation conditions to that of the patients, control participants were assigned to one of two conditions: they completed the task in either their left (n=14) or right (n=10) visual field.

The Institutional Review Boards of Carnegie Mellon University (CMU) and the University of Pittsburgh approved the study. All child participants provided informed assent, and their parents/guardians provided informed consent; adult participants consented for themselves. Participants received $25 as compensation.

### Experimental task

The two-rectangle paradigm used in this study (Figure 2) closely follows that of the original paradigm designed by Egly and colleagues^15^. We modified the task to make it more engaging and child-friendly, but the key parameters remain unchanged. As in the original task, a cue (a yellow outline) appeared in one of four locations on two objects, in this case cartoon ‘bones’ instead of rectangles, presented in the same, single visual field (left or right) across all trials. We also opted for a discrimination task in which participants identified, via key press, the color of the circle target ‘red’ or ‘purple’ which appeared at one end of a bone, as attentional cuing effects are consistently greater in discrimination tasks than in simple target detection tasks^35^.

**Figure 2.**
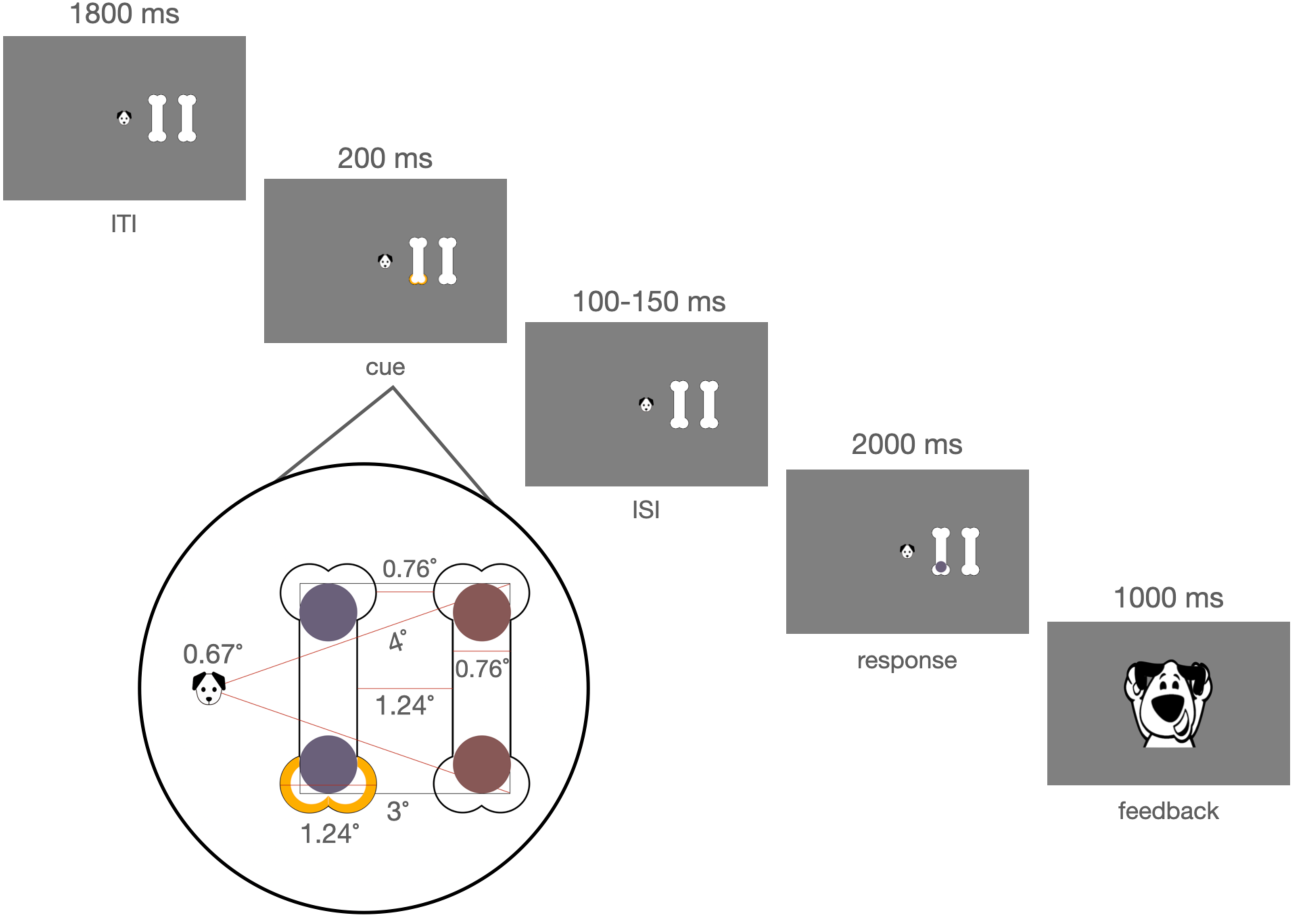
Two-rectangle attention paradigm modified to accommodate hemianopia in hemispheric surgery patients. Participants were instructed to fixate on ‘Doug the Dog’ and help him decide which colored ball (red or purple) appeared on his bones at any given time by pressing one key if they saw the red ball and another key if they saw the purple ball. They were not informed about the presence of the pre-cue that flashed in either a valid location (70% of trials), an invalid spatial location on the opposite object of the target to be presented (10% of trials, ‘IS’), an invalid location on the same object of the target to be presented (but equidistant to the cue to target distance of IS trials;10% of trials, ‘IO’), or in all locations (neutral trials: 10%). After the cue appeared and a jittered inter-stimulus interval (between 100-150 ms) passed, participants had 2000 ms to identify the color of the target. A happy ‘Doug’ was shown if a response was completed in the allotted time irrespective of accuracy, and a sad ‘Doug’ was shown if no response was completed.

Participants were instructed to have a ‘staring contest’ with the fixation marker, a small picture of a cartoon dog (’Doug the Dog’), while viewing the display and responding whether the color of Doug’s toy ball was red or purple as accurately and quickly as possible. The response keys to indicate ‘red’ and ‘purple’ were counter-balanced across participants. Fixation was monitored by the experimenter throughout the study. Also, because our interest is on covert rather than overt attention, we chose a brief exposure duration for the appearance of the cue and the ISI prior to target. It is unlikely, therefore, that the participants executed a saccade to the target as saccade preparation takes between 200-250 ms especially to unpredicted objects^40^. Last, post hoc analysis of controls’ performance between valid trials on the two bones showed an advantage in RT for the more foveally-versus more peripherally-located bones (z = -3.14, *p* = 0.002). This suggests that participants were likely centrally fixating rather than fixating between the two bones, as the latter viewing position would have equalized RTs across the two bones^41^. Note that we only used rectangles displayed vertically (and not horizontally) to reduce the length of the experiment and because Egly, Driver, et al.^15^ showed no significant difference of rectangle orientation (Experiment 1).

The timing of the stimulus presentation and details of stimulus sizes in visual degrees are described below and shown in Figure 2. A total of 400 experimental trials were pseudo-randomly split into 10 blocks (allowing the children to take breaks), preserving the 70:10:10:10 ratio of valid to IS to IO to neutral trials within blocks. Note that the number of IS and IO trials were the same. The maximum experiment time – if all 2000ms of allowed response time were used for each response – was 35 minutes.

### Stimulus Information

Stimuli were presented in PsychoPy^42^. Visual angles at which stimuli were presented are shown in Figure 2. The fixation marker, Doug the Dog, subtended 0.64 degrees vertically and horizontally. All stimuli were presented within approximately 4 degrees of the fixation marker, with two positions at approximately 1.8 degrees from fixation and the other two positions at approximately 3.6 degrees from fixation. The stimulus cue formed an outline of the edge of a bone and subtended 1.24 degrees. The target, a circle, subtended 0.74 degrees. Each bone had a minimum and maximum width of 0.74 and 1.24 degrees, respectively. The distance between an invalid cue and the target (either on the same object or at the same position on the other object) was approximately 3 degrees.

### Removal of neutral trials

We elected to use a neutral trial to assess the relative costs and benefits of valid and invalid trials. It became evident that the neutral trials, during which all four locations were cued, were not playing the intended role as a neutral, uninformative cue condition. Many children reacted with surprise when the four cues appeared and seemed uncertain how to proceed. As such, we decided to exclude these trials from further analysis.

### Software

All preprocessing and analysis were conducted in R^43^. For a list of all toolboxes used, see Table S2.

### Data Preprocessing

Three criteria were evaluated for data preprocessing. First, on a participant level, if the average accuracy was at or below chance (50%), that participant’s data were removed because such performance is indicative of a failure to perform the task. This step resulted in the removal of four patients (all LH surgery). Second, for the reaction time (RT) analyses, incorrect trials were removed for all participants. Third, trial counts for each trial type were calculated per participant. If a participant did not have at least 150 valid trials and 20 trials each of IS and IO, they were removed. For the accuracy analysis, which included incorrected trials, this step resulted in the removal of 1 LH and 1 RH surgery patient from the final accuracy dataset (16 LH and 13 RH surgery patients and 14 LH and 10 RH surgery matched controls). For the RT analysis, this step resulted in the removal of 1 control for whom stimuli were presented in their right field, 3 LH surgery patients, and 1 RH surgery patient. The RT dataset (final dataset: 14 LH and 13 RH surgery patients and 14 LH and 9 RH surgery matched controls) underwent a further preprocessing step in which RTs below the 5^th^ or above the 95^th^ percentile of a given participant’s RT distribution were replaced by the 5^th^ or 95^th^ percentile RT, respectively. This winsorization process reduces the impact of outliers while allowing for data retention^44^.

### Mixed Effects Modelling

#### Fixed Effects

We sought to examine how the within-subject fixed effect of trial type (valid, IS, or IO) varied across two between-subjects variables: the categorical but orthogonal factors of group (patients or controls) and hemifield of presentation (left or right).To maximize the power of the dataset, we performed analyses on trial-level observations across participants and fit random intercepts per participant to account for participant-level variability.

#### Age

Age was included as an additive fixed effect in all statistical models because we anticipated age affecting RTs^38,45,46^. Age was centered around the grand cohort mean so that the intercept was interpretable as the effect of the other predictors on response time for the average-aged participant. Age was not normalized or scaled so that the units of the effect could still be interpreted in terms of year.

#### Model Evaluation

Our linear mixed effects models (LMEMs) were fit to predict trial-level accuracy and RTs. The significance of each model term was evaluated with Type II Wald chi-squared analysis of deviance^47^ at an alpha criterion of 0.05. To determine the strength of evidence for each term in the model, a Bayes factor (BF), defined as the ratio of the prior predictive probabilities, was approximated using the Bayesian Information Criteria (BIC) for the null model that does not include the term subtracted from the BIC for the alternative model with the term^48^. A BF greater than 3 therefore offers evidence for the null hypothesis, while a BF less than 0.33 offers evidence for the alternative hypothesis^49^. Post-hoc comparisons at each level of the significant factors were then calculated and corrected for multiple comparisons using a false discovery rate of < 0.05^50^.

## Results

To evaluate spatial- and object-based attention within a single hemisphere, we compared patients who underwent childhood hemispheric surgery to TD age-matched controls on the paradigm established by Egly, Driver, et al.^15^, modified here for appropriateness for children (see Figures 1 and 2). Overall, participants performed the task with high accuracy: on average, controls responded correctly in 94% of trials and patients responded correctly in 85% of trials, where chance was 50% (red/purple ball). Controls had significantly higher accuracy than patients (z = 3.12, *p* = 0.002), but this group difference did not interact with trial type or any other variable of interest (see Table S3 for full description of accuracy results).

For our main dependent variable of interest, RT, only correct trials were analysed to ensure that patients were in fact attending to the target when they responded. Participants’ average RTs for correct responses to each of the three trial types (valid, IS, and IO) and the average RT costs for each type of invalid cuing (relative to valid, calculated by subtracting each participant’s average RT for the valid cue condition from that for each invalid cue type), are shown in Figure 3.

**Figure 3.**
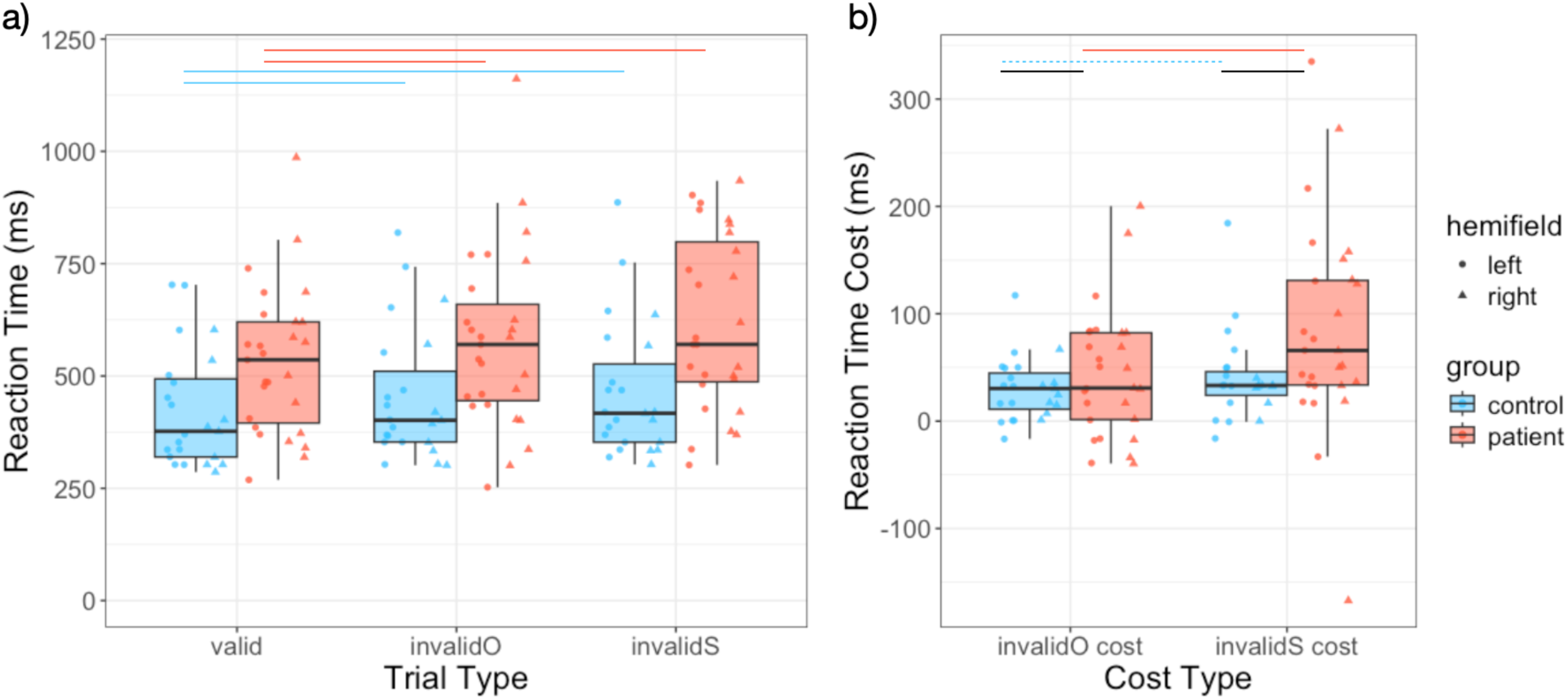
Distribution of (a) reaction times and (b) invalid cue costs for patients (red) and controls (blue) for each trial type and cost type, respectively. The line in each box represents the median, while the lower and upper ranges represent the 25^th^ and 75^th^ percentiles, respectively; the whiskers extend to 1.5 times the interquartile range. Reaction time costs were calculated for each participant by subtracting their median RT on valid trials from that on invalid object trials (invalidO cost) and their median RT on valid trials from that on invalid spatial trials (invalidS cost). The lines above the boxplots represent post-hoc comparisons that were tested between trial/cost types within controls (blue) or within patients (red) and between groups within cost type (black). Solid lines represent significant differences and dashed lines represent differences that were tested but were not significant. As stimuli were only presented to patients’ intact hemifield, i.e., the hemifield opposite to their intact hemisphere. Thus, LH surgery patients saw stimuli in their left hemifield (circles) and RH surgery patients saw stimuli in their right hemifield (triangles). Statistical details are included in the main text.

### Model 1: Comparison of controls and patients for spatial- and object-based attention

In the first linear mixed effects model (LMEM), we modelled RT from the three-way interaction between group (patients, controls), hemifield of presentation (left, right), and trial type (valid, IO, IS), with mean-centered age as an additive effect and participant as a random effect. The three-way interaction term was not significant (group x hemifield x trial type: X^2^(2, *n* = 50) =0.817, *p* = 0.665, BF = 10633.42). There was also no significant effect of hemifield as a function of group (group x hemifield: X^2^(1, *n* = 50) = 1.13, *p* = 0.289, BF = 72.49) or of trial type (trial type x hemifield: X^2^(2, *n* = 50) = 1.65, *p* = 0.438, BF = 7010.24). However, patients’ and controls’ RTs were differentially affected by trial type (group x trial type: X^2^(2, *n* = 50) = 20.83, *p* < 0.0001, BF = 0.48), as detailed in the post-hoc comparisons below. Beyond this interaction, we also found significant main effects of group (X^2^(1, *n* = 50) = 12.23, *p* = 0.0005, BF = 0.59), trial type (X^2^(2,*n* = 50) = 247.56, *p* < 0.0001, BF < 0.0001), and age (X^2^(1, *n* = 50) = 14.11, *p* = 0.0002, BF = 0.23). Older participants responded faster than younger participants, with an estimated 12 ms increase per yr of age (t = -3.76, *p* = 0.0005), and also had lower invalid trial costs (see Figure S1). There was no significant main effect of hemifield (X^2^(1, *n* = 50) = 0.174, *p* = 0.676, BF = 116.22) on RT.

Close scrutiny of the costs in RT (Figure 3b) revealed that at least 4 participants had very high costs, at above 200 ms or below -100 ms. We recomputed the statistics while excluding these individuals, who were all patients, to ensure that the outcome was not simply a result of these potential outliers. The results were unchanged. Together, the re-analysis confirms the robustness of the longer IS than IO cost in the patients over controls.

#### Analysis of the validity effect

Within group post hoc comparisons of the group x trial type interaction revealed that, although patients responded more slowly than controls (z = 3.94, *p* = 0.0001), both patients and controls showed a significant validity effect in which RTs to the valid condition were significantly faster than to the IO and IS conditions (control IO cost: 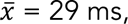 z = 4.80, *p* < 0.0001; control IS cost: 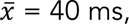 z = 6.33, *p* < 0.0001; patient IO cost: 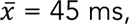 z = 8.50, *p* < 0.0001; patient IS cost: 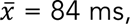 z = 12.45, *p* < 0.0001). This result is important in demonstrating that patients can take advantage of the cue and that this is so regardless of which hemisphere was resected.

#### Comparison of IS and IO trials

Post hoc comparisons of IS and IO conditions revealed that, contrary to the adult literature^15^, controls in our largely pediatric sample did not show a significant RT difference between IS and IO trials (Δ = 11 ms, z = 1.18, *p* = 0.237). In contrast, patients did have significantly longer RTs on IS trials relative to IO trials (Δ = 39 ms, z = 4.65, *p* < 0.0001). We also compared invalid trials across groups by subtracting each group’s estimated marginal mean of the valid trials from their IO and IS trials. Patients had significantly greater costs relative to controls for both IO trials (Δ = 16 ms, z = 2.32, *p* = 0.026) and IS trials (Δ = 44 ms, z = 3.45, *p* = 0.0009). Together, these results suggest that the difference in RTs for IS and IO trials for patients is more likely a result of the increased cost on IS trials and less likely a result of an advantage on IO trials, as controls did not show a significant advantage for IO compared to IS trials.

### Model 2: Left and right hemispheric surgery patients are equivalently impaired in object- and spatial-based attention

Because we had a clear set of a priori predictions of hemispheric differences in the patients, based on Egly, Driver, et al.^51^, we fit a second LMEM using only the patient data, and modelled the prediction of RT from preserved hemisphere (left, right) and trial type (valid, IO, IS). There was no significant interaction of preserved hemisphere x trial type on RT (X^2^(2, *n* = 27) = 0.211, *p* = 0.900, BF = 7372.08). Importantly, the main effect of preserved hemisphere was also not significant (X^2^(1, *n* = 27) = 0.627, *p* = 0.429, BF = 66.41). There was, however, as above, a significant main effect of trial type on RT (X^2^(2, *n* = 27) = 138.14, *p* < 0.0001, BF < 0.0001). Post-hoc comparisons on trial type confirmed that responses to the valid condition were significantly faster than those to the IO and IS conditions (IO cost: z = 6.97, *p* < 0.0001; IS cost: z = 10.22, *p* < 0.0001), and that the IS cost was significantly greater than the IO cost (z = -2.83, *p* = 0.005), irrespective of hemisphere preserved.

### Summary

Together, these analyses reveal that, unsurprisingly, patients performed more slowly than controls (group main effect), and RTs increased with age (age main effect). We also documented the expected profile across trial types^15^, with both patients and controls performing faster on valid than on invalid trials, demonstrating the advantage of the attentional cue in both groups. For the patients, this was true irrespective of hemisphere resected, and for controls, there was no effect of hemifield of presentation. Intriguingly, our (largely pediatric) controls do not show the difference in RT between IO vs IS trials that has been previously reported in adults and this held equally over visual field of presentation^15^. Additionally, there was an unexpected modulation of trial type by group: patients’ RTs on IO and IS trials were significantly longer than those of controls, and within patients, there were longer RTs for IS trials than IO trials. This finding suggests that neither group demonstrates an object-based advantage conferred on performance when targets are shown on a cued object, and patients have a larger cost in switching between locations on different objects than within the same object. Patients’ object-switching cost was also independent of which hemisphere was preserved during development.

## Discussion

The current study explored spatial- and object-based attention in individuals who have undergone childhood hemispheric surgery, providing novel insights into the effects of confining the development of visual attentional processes to a single hemisphere. There were two key findings. The first was that, surprisingly and interestingly, the control group did not show the benefit that typically accrues for targets in locations on the cued object compared to the uncued object, as was robustly demonstrated in adults^15^. Second, patients who develop with just a single hemisphere, be it left or right, maintain the facilitative advantage afforded by valid cuing, but are significantly more delayed by invalid cues to targets either within the cued object (IO) or in the uncued object (IS) compared to controls. Furthermore, patients had an addition cost of switching attention across objects (IS) relative to within object (IO) that was not observed in our controls.

### Controls show equivalent performance on object- and space-based trials

It is interesting that the control group, largely comprised of children, do not show the advantageous modulation of spatial distance when the invalid target is located in the cued versus uncued object. These results indicate that the hemispheric lateralization of object and space-based attention may evolve over the course of development, a proposal that remains to be verified with cross-section across age or longitudinal data. Changes in visual search and the increase in RH participation in complex visual search is compatible with this hypothesis^26^. There are other findings, albeit not in the domain of attentional processing, that report evolving changes from bilateral to more unilateral organization over development. For example, in the domain of language, the left and right hemispheres appear to be equipotential early in development but then, in most individuals, become more LH dominant with age, although a ‘shadow’ remains in the RH^52^. Likewise, early in development, unlike in adulthood, both hemispheres appear to subserve the representation of faces and damage to either hemisphere results in a recognition deficit^53^. Related research reveals that once literacy is acquired, word processing appears to become more LH dominant^54^, perhaps to be in close proximity to LH language areas. Similarly, face processing becomes more lateralized to the RH, perhaps to be in closer proximity to regions associated with social behaviors^55^. The emerging picture is of hemispheric functions being in flux as the multiple cognitive abilities and their neural substrates are configured over the course of development. Indeed, the evolving lateralization in multiple cognitive and social domains simultaneously constitutes an interesting constraint-satisfaction problem and a broad investigation of lateralization tracking the brain-behavior changes over multiple processing domains would be instructive in understanding how the mature adult-like lateralization profile emerges.

### Patients exhibit a validity effect and significantly larger IS compared to IO costs

Hemispheric surgery patients exhibited a robust validity effect, demonstrating intact sensitivity to attentional cues. Unsurprisingly, patients were slower to respond compared to controls, perhaps a consequence of impaired motor control in hemispheric surgery patients^39,56^. The fact that patients nonetheless responded faster on valid vs invalid trials demonstrates that mechanisms that underlie the facilitation of a highly predictive cue in enhancing processing at a spatial location can remain operational despite the loss of an entire cerebral hemisphere, either left or right, during early development. This is consistent with previous findings suggesting that certain cognitive functions can be largely preserved when a single hemisphere develops in isolation^38^. On the other hand, we observed that patients exhibited significantly larger costs for IS compared to IO trials, and, compared to controls, had significantly larger costs for both types of invalid targets, in IO trials (IO cost: 45 ms for patients, 29 ms for controls) and to a greater extent in IS trials (IS cost: 84 ms for patients, 40 ms for controls). As controls did not show this longer latency for IS trials, these results suggest that development of space and object-based attention in a single hemisphere might negatively affect the ability to efficiently switch attention across objects.

### Side of resection does not predict patients’ overall RTs or RTs to different trial types

Given the established lateralization of space- and object-based attention to the respective RH and LH in non-neurological adults^13,22^, it is notable that patients’ performance was not affected by which hemisphere was resected in RT or accuracy overall or as a function of trial type^31,though see:, 32^. In particular, these patient findings are comparable to results observed by Egly, Rafal, and colleagues ^51^ specifically in the LH of the split-brain patient JW. Patient JW had similar RTs for IO and IS trials presented to the RH and comparable RTs to IO trials presented to the RH and LH, but slower RTs for IS trials presented to the LH. While both hemispheres might be equivalent for space- and object-based attention in early development, and damage to either hemisphere produces the same outcome, in adulthood when the attentional processes are lateralized, the LH plays the more critical role in object-based attention than the RH.

In addition to their research with the split-brain patient, Egly, Driver, and colleagues^51^ also acquired data using a similar paradigm from another neuropsychological population: patients with either LH or RH parietal damage (two of whom have a circumscribed parietal resection). This study showed that LH parietal damage patients have a sensitivity to object-related shifts of attention that RH damaged patients did not. However, they presented one rectangle in each visual field in both horizontal and vertical orientations, and then collapsed across the orientation. This resulted in IO and IS cue-target pairs that were presented within the same hemifield, as in the IO condition in our study, to be collapsed with IO and IS pairs presented across hemifields, which was not possible in our study due to hemianopia in our patients. Because Egly, Driver, and colleagues^15^ only analysed LH and RH parietal patients for trial type differences based on the hemifield of the target (collapsing across orientation and, therefore, collapsing within and across hemifield shifts), their study differs sufficiently from ours to make further comparisons difficult.

### Potential mechanisms for the atypical patients’ attention profile

Two patient findings require explanation: the lack of an effect of which hemisphere is resected and the larger IS than IO cost that was not observed in controls. Below, we offer potential accounts for each of these results.

#### Lack of hemisphere difference

First, the timing of lateralization of object- and space-based attention in the typically developing population is poorly understood, with some studies suggesting that the characteristic right-lateralization of spatial attention observed in adults emerges around age 10 and others demonstrating spatial functions that remain consistently bilateral (e.g.^13,24,26^). It is thus possible that the age at which the children underwent hemispheric surgery in this study (median age = 14.89 yr) was before the maturation of lateralized attentional systems. It is also possible that, even though there was no effect of hemifield of presentation in this study, controls might have some degree of hemispheric lateralization that could not be behaviorally demonstrated because information is shared across both hemispheres after input, masking the effect of presenting to only one hemifield. If this is the case, and if our patients also had hemispheric differentiation *before* surgery, it possible that *after* surgery, the LH and RH might be equally maladaptively affected by brain injury. For example, perhaps as a result of ‘neural crowding’^57^, the more spatially biased RH might struggle to accommodate object-based processing after LH resection and the more object biased LH might struggle to accommodate spatial-based processing after RH resection, leading to the lack of hemisphere differences observed here.

#### Larger IS than IO cost

On the face of it, it is surprising that the patients have longer reaction times to the uncued object when controls do not. This failure in switching to the uncued object or ‘object-based disengagement’, akin to Posner’s disengage space-based deficit in hemispatial neglect requires explanation. One possibility is that, in bottom-up fashion, the patients fail to perceptually organize the scene into two rectangles – it is only the arrival of the cue and the subsequent automatic attention spread within the bounds of the cued rectangle^58^ that both facilitate the valid targets and the targets in the cued rectangle. On this account, the IS cost derives from the fact that the uncued object remains unparsed. An alternative account posits that the rectangles have both been parsed (i.e., the scene has been segmented and perceptually organized) and the target at the cued location and within the cued object both benefit from the cueing. The greater cost when switching attention to a location on a different object might then reflect a deficit in the ability to disengage object-based attention from the cued object – because the valid trials and IO trials constitute 80% of all trials (10% IS and 10% neutral), the cued rectangle is assigned high prioritization and the high statistical probability inhibits the disengagement. These two accounts may not be mutually exclusive and cannot be adjudicated between based on the data reported here. However, attempts have been made to separate these two possibilities. In typical observers, Ho^59^ varied the load (high versus low) and showed that, only on the high (color/shape conjunction), but not the low (color feature), load condition, was the benefit of the invalid target in the cued rectangle observed, thereby favoring the spread of attention in the cued object or the sensory enhancement associated with the clue. It is the case, however, that other studies that directly pit these accounts against each other reveal that depending on task contingencies, strength of the object representation, and timing, the cued versus uncued object advantage can be altered^60,61^.

### Limitations

A potential limitation of our study was that, to limit the overall length of the study so that children would be able to complete the task, we did not evaluate both vertically and horizontally oriented objects (bones). As such, in the IO condition, the attention shifted vertically from the cue to the opposite location on the same object, whereas shifts across objects in the IS condition always involved the horizontal direction. Though Egly, Driver, and colleagues^15^ did not find a significant effect of object orientation when presenting horizontally compared to vertically, we cannot entirely rule out that the difference between our patients and controls in trial type is a specific deficit in horizontal (IS) versus vertical (IO) shifts in attention. Another limitation is the inability to determine valid and invalid costs and benefits relative to a baseline: if the neutral condition in our experiment had served as a good uninformative cue, and not as a distracting surprise, we would have been able to distinguish the facilitative effects of the valid cue from the detrimental effects of the invalid cues. However, given children’s abnormal responses to these cues, invalid cuing costs in our results were inherently tied to valid cuing benefits.

Yet a further limitation of this study is the heterogeneity of our sample, with participants spanning a wide age range and undergoing hemispheric surgery at different ages. While we accounted for age as a fixed effect in our analyses (and, indeed, it is statistically significant), future studies with a greater and more balanced sampling of ages should investigate attentional processing in controls and patients to better understand how the timing of surgery and brain development interact to influence attentional capacities. Additionally, etiology of disease (e.g., dysplasia vs stroke) may have notable independent effects on postoperative cognitive outcomes, such as attention^62,63^, which we cannot capture with our relatively small sample. Lastly, it is important to acknowledge that attention is a broad cognitive domain, with many underlying mechanisms; only a specific attentional trade-off between spatial- and object-based attention was explored here. There are also other forms of attention, for example, attention to particular features of the input^6^, but these types of attention should be explored in hemispheric surgery patients in future studies.

Last, but not least, there has been some controversy regarding the two-rectangle paradigm. Among the criticisms is the claim that this paradigm cannot adjudicate whether the object-based advantage derives from a truly object-based representation that codes for object structure or from a representation in which spatial locations are grouped. If the latter holds, then the results have implications for space-based processing and object-based processing is irrelevant^64^. A second concern relates to the potential hemispheric differences, especially the claim that object-based attention is mediated by the LH. Valsangkar-Smyth et al.^65^, using a modification of the object-based attention paradigm designed by Duncan^66^, showed a greater object cost when the visual displays were in the right visual field, i.e., the LH, compared to when they were shown in the left visual field, i.e. the RH. Relatedly, using the same paradigm as Valsangkar-Smyth et al.^65^ with the same split-brain patient as Egly, Rafal et al.^51^, Kingstone^67^ also reported that object-based attention is lateralized to the RH (evident in the left but not right hemifield).

Although the specifics of the debate about the two-rectangle paradigm do not impugn the current results (as we examined a difference between two groups on the same paradigm), it is worth noting that the paradigms used may result in different outcomes and the debate is not settled.

## Conclusion

The current study characterizes spatial- and object-based attentional processing in typically developing controls and patients who underwent early hemispheric surgery. The findings suggest that patients with a single hemisphere can benefit from valid cues and, hence, maintain efficient attentional processing similar to that of typically developing controls, but have significantly longer latencies for invalidly cued targets compared to controls. The increased cost associated with object-switching in these patients indicates that the attentional system, when constrained to a single hemisphere, results in a disproportionate cost of shifting attention to a location on an uncued object than on the cued object. That our LH and RH patients had an indistinguishable profile of difficulty with invalidly cued locations on cued and uncued objects could be explained by 1) malleability in the lateralization of spatial- and object-based attention or maladaptive plasticity that affects lateralized LH and RH functions in the same way, or 2) a bilateral distribution of spatial- and object-based attention at the stage of development during which they underwent surgery. Future research should explore the trajectory of space- and object-based attentional processing over development to ask how the timing of surgery in childhood resection might affect the lateralization of attentional systems.

## Competing interest statement

Behrmann is a co-founder of and holds equity in Precision Neuroscopics. The other authors declare no competing financial interests.

## Acknowledgments

S.R. was supported by the National Science Foundation Graduate Research Fellowship Program, Grant No. DGE2140739. M.C.G. was supported by T32GM144300 from the National Institute of General Medical Sciences and a scholarship from the University of Pittsburgh MD-PhD program. The research was also supported by the National Eye Institute grant (R01 EY027018) to M.B. who also acknowledges support from P30 CORE award EY08098 from the NEI, as well as unrestricted supporting funds from The Research to Prevent Blindness Inc, NY and the Eye & Ear Foundation of Pittsburgh. The content is solely the responsibility of the authors and does not necessarily represent the official views of the NEI, NIGMS, NSF, or the University of Pittsburgh. We thank the Pediatric Epilepsy Surgery Alliance members, its Chief Executive Officer, Monika Jones, and the many participants and families who have given generously their time for this study.

## Data and Code Availability

All raw data, as well as code for the experimental task and preprocessing/analysis, will be made freely available upon publication on the CMU KiltHub repository (reserved digital object identifier: 10.1184/R1/27221211).

## References

1. Eckstein, M. P. Probabilistic Computations for Attention, Eye Movements, and Search. Annu. Rev. Vis. Sci. 3, 319–342 (2017).

2. Wolfe, J. M. Visual Search: How Do We Find What We Are Looking For? Annu. Rev. Vis. Sci. 6, 539–562 (2020).

3. Corbetta, M., Kincade, J. M., Ollinger, J. M., McAvoy, M. P. & Shulman, G. L. Voluntary orienting is dissociated from target detection in human posterior parietal cortex. Nat. Neurosci. 3, 292–297 (2000).

4. Geng, J. J. & Behrmann, M. Spatial probability as an attentional cue in visual search. Percept. Psychophys. 67, 1252–1268 (2005).

5. Posner, M. I. Orienting of attention. Q. J. Exp. Psychol. 32, 3–25 (1980).

6. Posner, M. I., Walker, J. A., Friedrich, F. J. & Rafal, R. D. Effects of parietal injury on covert orienting of attention. J. Neurosci. 4, 1863–1874 (1984).

7. Bartolomeo, P. & Seidel Malkinson, T. Hemispheric lateralization of attention processes in the human brain. Curr. Opin. Psychol. 29, 90–96 (2019).

8. Gaffan, D. & Hornak, J. Visual neglect in the monkey. Representation and disconnection. Brain J. Neurol. 120, 1647–1657 (1997).

9. Cai, Q., Van der Haegen, L. & Brysbaert, M. Complementary hemispheric specialization for language production and visuospatial attention. Proc. Natl. Acad. Sci. 110, E322–E330 (2013).

10. Karnath, H. O., Ferber, S. & Himmelbach, M. Spatial awareness is a function of the temporal not the posterior parietal lobe. Nature 411, 950–953 (2001).

11. Corbetta, M., Miezin, F. M., Dobmeyer, S., Shulman, G. L. & Petersen, S. E. Attentional Modulation of Neural Processing of Shape, Color, and Velocity in Humans. Science 248, 1556–1559 (1990).

12. Corbetta, M. et al. A Common Network of Functional Areas for Attention and Eye Movements. Neuron 21, 761–773 (1998).

13. Scolari, M., Seidl-Rathkopf, K. N. & Kastner, S. Functions of the human frontoparietal attention network: Evidence from neuroimaging. Curr. Opin. Behav. Sci. 1, 32–39 (2015).

14. Coll, S. Y., Marti, E., Doganci, N. & Ptak, R. The disengagement deficit after right-hemisphere damage: Distinct roles of lateral frontal and parietal damage. Brain Res. Bull. 214, 111003 (2024).

15. Egly, R., Driver, J. & Rafal, R. D. Shifting visual attention between objects and locations: Evidence from normal and parietal lesion subjects. J. Exp. Psychol. Gen. 123, 161–177 (1994).

16. Li, K. & Malhotra, P. A. Spatial neglect. Pract. Neurol. 15, 333–339 (2015).

17. Behrmann, M. Spatial neglect: Cognitive and neural bases. Brain Cogn. 2, 194–195 (2007).

18. Machner, B. et al. Behavioral deficits in left hemispatial neglect are related to a reduction of spontaneous neuronal activity in the right superior parietal lobule. Neuropsychologia 138, 107356 (2020).

19. Whitwell, R. L., Striemer, C. L., Cant, J. S. & Enns, J. T. The Ties that Bind: Agnosia, Neglect and Selective Attention to Visual Scale. Curr. Neurol. Neurosci. Rep. 21, 54 (2021).

20. Fiebelkorn, I. C. & Kastner, S. Functional Specialization in the Attention Network. Annu. Rev. Psychol. 71, 221–249 (2020).

21. Hollingworth, A., Maxcey-Richard, A. M. & Vecera, S. P. The spatial distribution of attention within and across objects. J. Exp. Psychol. Hum. Percept. Perform. 38, 135–151 (2012).

22. Shomstein, S. & Behrmann, M. Cortical systems mediating visual attention to both objects and spatial locations. Proc. Natl. Acad. Sci. 103, 11387–11392 (2006).

23. Ebner, K., Lidzba, K., Hauser, T.-K. & Wilke, M. Assessing language and visuospatial functions with one task: A “dual use” approach to performing fMRI in children. NeuroImage 58, 923–929 (2011).

24. Ferrara, K., Seydell-Greenwald, A., Chambers, C. E., Newport, E. L. & Landau, B. Developmental changes in neural lateralization for visual-spatial function: Evidence from a line-bisection task. Dev. Sci. 25, e13217 (2022).

25. Everts, R. et al. Strengthening of laterality of verbal and visuospatial functions during childhood and adolescence. Hum. Brain Mapp. 30, 473–483 (2009).

26. Lidzba, K., Ebner, K., Hauser, T.-K. & Wilke, M. Complex Visual Search in Children and Adolescents: Effects of Age and Performance on fMRI Activation. PLOS ONE 8, e85168 (2013).

27. Johnson, M. H., Posner, M. I. & Rothbart, M. K. Facilitation of Saccades Toward a Covertly Attended Location in Early Infancy. Psychol. Sci. 5, 90–93 (1994).

28. Ross-Sheehy, S., Schneegans, S. & Spencer, J. P. The Infant Orienting With Attention task: Assessing the neural basis of spatial attention in infancy. Infancy OI. J. Int. Soc. Infant Stud. 20, 467–506 (2015).

29. Schatz, J., Craft, S., Koby, M. & DeBaun, M. R. A lesion analysis of visual orienting performance in children with cerebral vascular injury. Dev. Neuropsychol. 17, 49–61 (2000).

30. Smith, S. E. & Chatterjee, A. Visuospatial attention in children. Arch. Neurol. 65, 1284–1288 (2008).

31. Trauner, D. A. Hemispatial neglect in young children with early unilateral brain damage. Dev. Med. Child Neurol. 45, 160–166 (2003).

32. Hart, E. et al. Pediatric unilateral spatial neglect: A systematic review. J. Pediatr. Rehabil. Med. 14, 345–359 (2021).

33. Marsh, E. B. et al. Hemispherectomy sustained before adulthood does not cause persistent hemispatial neglect. Cortex J. Devoted Study Nerv. Syst. Behav. 45, 677–685 (2009).

34. Adamos, T., Chukoskie, L., Townsend, J. & Trauner, D. Spatial attention in children with perinatal stroke. Behav. Brain Res. 417, 113614 (2022).

35. Kinsbourne, M. Orientational bias model of unilateral neglect:Evidence from attentional gradients within hemispace. Unilateral Negl. Clin. Exp. Stud. Hove (1993).

36. Danguecan, A. N. & Smith, M. L. Re-examining the crowding hypothesis in pediatric epilepsy. Epilepsy Behav. EB 94, 281–287 (2019).

37. Dennis, M. et al. Functional Plasticity in Childhood Brain Disorders: When, What, How, and Whom to Assess. Neuropsychol. Rev. 24, 389 (2014).

38. Granovetter, M. C., Robert, S., Ettensohn, L. & Behrmann, M. With childhood hemispherectomy, one hemisphere can support—but is suboptimal for—word and face recognition. Proc. Natl. Acad. Sci. 119, e2212936119 (2022).

39. Kim, J., Park, E.-K., Shim, K.-W. & Kim, D. S. Hemispherotomy and Functional Hemispherectomy: Indications and Outcomes. J. Epilepsy Res. 8, 1–5 (2018).

40. Yang, J. et al. Analysis of Predictive Factors in Surgical Treatment of Intractable Epilepsy Caused by Focal Cortical Dysplasia in Children. Int. J. Neurosci. 0, 1–11 (2024).

41. Downing, C. J. & Pinker, S. The Spatial Structure of Visual Attentio. in Attention and Performance XI (Routledge, 1985).

42. Peirce, J. et al. PsychoPy2: Experiments in behavior made easy. Behav. Res. Methods 51, 195–203 (2019).

43. R Core Team. R: A Language and Environment for Statistical Computing. (R Foundation for Statistical Computing, Vienna, Austria, 2020).

44. Wilcox, R. R. 3 - SUMMARIZING DATA. in Applying Contemporary Statistical Techniques (ed. Wilcox, R. R.) 55–91 (Academic Press, Burlington, 2003). doi:10.1016/B978-012751541-0/50024-9.

45. Bucsuházy, K. & Semela, M. Case Study: Reaction Time of Children According to Age. Procedia Eng. 187, 408–413 (2017).

46. Kiselev, S., Espy, K. A. & Sheffield, T. Age-related differences in reaction time task performance in young children. J. Exp. Child Psychol. 102, 150–166 (2009).

47. Fox, J. & Weisberg, S. An R Companion to Applied Regression. (SAGE Publications, 2018).

48. Wagenmakers, E.-J. A practical solution to the pervasive problems ofp values. Psychon. Bull. Rev. 14, 779–804 (2007).

49. Lee, M. & Wagenmakers, E.-J. Bayesian Cognitive Modeling : A Practical Course. Bayesian Cogn. Model. Pract. Course 265 (2013) doi:10.1017/CBO9781139087759.

50. Benjamini, Y. & Yekutieli, D. The control of the false discovery rate in multiple testing under dependency. Ann. Stat. 29, 1165–1188 (2001).

51. Egly, R., Rafal, R., Driver, J. & Starrveveld, Y. Covert Orienting in the Split Brain Reveals Hemispheric Specialization for Object-Based Attention. Psychol. Sci. 5, 380–383 (1994).

52. Martin, K. C. et al. A Weak Shadow of Early Life Language Processing Persists in the Right Hemisphere of the Mature Brain. Neurobiol. Lang. Camb. Mass 3, 364 (2022).

53. de Schonen, S., Mancini, J., Camps, R., Maes, E. & Laurent, A. Early brain lesions and face-processing development. Dev. Psychobiol. 46, 184–208 (2005).

54. Dundas, E. M., Plaut, D. C. & Behrmann, M. Variable left-hemisphere language and orthographic lateralization reduces right-hemisphere face lateralization. J. Cogn. Neurosci. 27, 913–925 (2015).

55. Blauch, N. M., Plaut, D. C., Vin, R. & Behrmann, M. Individual variation in the functional lateralization of human ventral temporal cortex: Local competition and long-range coupling. 2024.10.15.618268 Preprint at 10.1101/2024.10.15.618268 (2024).

56. van Empelen, R., Jennekens-Schinkel, A., Buskens, E., Helders, P. J. M. & van Nieuwenhuizen, O. Functional consequences of hemispherectomy. Brain 127, 2071–2079 (2004).

57. Lidzba, K., Staudt, M., Wilke, M. & Krägeloh-Mann, I. Visuospatial deficits in patients with early left-hemispheric lesions and functional reorganization of language: consequence of lesion or reorganization? Neuropsychologia 44, 1088–1094 (2006).

58. Ekman, M., Roelfsema, P. R. & Lange, F. P. de. Object Selection by Automatic Spreading of Top-Down Attentional Signals in V1. J. Neurosci. 40, 9250–9259 (2020).

59. Ho, M.-C. Object-Based Attention: A Within-Object Benefit and Sensory Enhancement in Discrimination Tasks. 中華心理學刊 55, 181–199 (2013).

60. Shomstein, S. & Behrmann, M. Object-based attention: Strength of object representation and attentional guidance. Percept. Psychophys. 70, 132 (2008).

61. Shomstein, S. & Yantis, S. Configural and contextual prioritization in object-based attention. Psychon. Bull. Rev. 11, 247–253 (2004).

62. Jehi, L. & Braun, K. Does etiology really matter for epilepsy surgery outcome? Brain Pathol. 31, e12965 (2021).

63. Pulsifer, M. B. et al. The cognitive outcome of hemispherectomy in 71 children. Epilepsia 45, 243–254 (2004).

64. Vecera, S. P. Grouped locations and object-based attention: Comment on Egly, Driver, and Rafal (1994). J. Exp. Psychol. Gen. 123, 316–320 (1994).

65. Valsangkar-Smyth, M. A., Donovan, C.-L., Sinnett, S., Dawson, M. R. W. & Kingstone, A. Hemispheric performance in object-based attention. Psychon. Bull. Rev. 11, 84–91 (2004).

66. Duncan, J. Selective attention and the organization of visual information. J. Exp. Psychol. Gen. 113, 501–517 (1984).

67. Kingstone, A. Covert orienting in the split brain: Right hemisphere specialization for object-based attention. Laterality 21, 732–744 (2016).

