## Supplemental Information for "Space- and object-based attention in patients with a single hemisphere following childhood resection"

**Table S1**. Biographical and medical information on patient population.

| **Age** | **Gender** | **~Age at Seizure Onset (yr)** | **~Age at First Surgery (yr)*** | **~Age at Last Surgery (yr)*** | **Cognitive Ability**** |
| --- | --- | --- | --- | --- | --- |
| **Left Hemisphere Surgery Cases** | | |  |  | |
| 13.98 | Male | 11.00 | 15.00 | 16.00 | - |
| 14.87 | Female | 0.33 | 0.75 | 10.92 | mildy impaired |
| 14.51 | Male | 0.42 | 4.77 | | mild to moderately impaired |
| 11.37 | Female | 0.17 | 5.53 | 10.78 | slightly delayed |
| 20.28 | Female | 1.67 | 4.86 | | mild impairment |
| 13.12 | Female | 0.17 | 0.53 | 1.37 | mildly impaired |
| 19.21 | Male | 4.00 | 13 | 14 | moderately impaired |
| 14.21 | Female | 1.17 | 3.06 | | moderately impaired |
| 25.28 | Male | 0.00 | 8 | | above average |
| 16.54 | Female | 0.50 | 3.89 | 4.97 | moderately impaired |
| 15.47 | Female | 0.00 | 1.08 | | moderately impaired |
| 30.08 | Female | 0.00 | 6.23 | | above average |
| 5.95 | Female | 1.08 | 3.06 | | average |
| 12.30 | Female | 1.50 | 10 | | mild to moderately impaired |
| 8.90 | Female | 0.00 | 0.84 | | delayed |
| 14.35 | Male | 2.00 | 7.91 | | moderately impaired |
| 5.53 | Male | 0.00 | 0.35 | | mildly impaired |
| 7.31 | Female | 3.00 | 4.24 | | average |
| 13.78 | Male | 3.00 | 6.68 | 14.3 | average |
| 12.50 | Female | - | - | | - |
| 14.55 | Male | - | - | | - |
| **Right Hemisphere Surgery Cases** | | |  |  | |
| 20.77 | Female | 0.00 | 9.24 | | average |
| 16.18 | Female | 1.00 | 1.30 | | average |
| 13.24 | Female | 0.00 | 0.61 | 9.83 | average |
| 12.12 | Female | 0.25 | 0.65 | 2.31 | mil y impaired |
| 32.41 | Female | 5.00 | 8.52 | | moderately impaired |
| 14.90 | Female | 2.00 | 6.09 | | moderately impaired |
| 19.35 | Male | 0.25 | 6.64 | | significantly impaired |
| 17.03 | Male | 0.00 | 0.98 | | mildly impaired |
| 9.14 | Male | 0.42 | 3.42 | 7.42 | moderately impaired |
| 18.01 | Male | 0.58 | 0.00 | 3.06 | mildly impaired |
| 5.06 | Male | 0.13 | 0.09 | 3.59 | average |
| 20.68 | Male | 0.08 | 2.43 | 7.01 | mildly impaired |
| 19.19 | Female | 4.00 | 6.42 | | mildly impaired |
| 20.04 | Female | 3.00 | 3.37 | | mildly impaired |

*Surgery and seizure onset ages are approximate and per participant and/or guardian report. Some patients had surgeries before complete hemispheric surgeries, e.g., a more focal resection or ablation before hemispherectomy.

** Per participant or guardian report.

- No information available.

**Table S2**. List of R (version 4.4.1) packages and toolboxes used for analyses.

| **Package** | **Version** |
| --- | --- |
| gridExtra | 2.3 |
| interactions | 1.2.0 |
| performance | 0.12.3 |
| ggsignif | 0.6.4 |
| broom | 1.0.6 |
| psych | 2.4.6.26 |
| emmeans | 1.10.4 |
| lmerTest | 3.1-3 |
| lme4 | 1.1-35.5 |
| Matrix | 1.7-0 |
| plyr | 1.8.9 |
| pracma | 2.4.4 |
| rstudioapi | 0.16.0 |
| lubridate | 1.9.3 |
| forcats | 1.0.0 |
| stringr | 1.5.1 |
| dplyr | 1.1.4 |
| purrr | 1.0.2 |
| readr | 2.1.5 |
| tidyr | 1.3.1 |
| tibble | 3.2.1 |
| ggplot2 | 3.5.1 |
| tidyverse | 2.0.0 |
| car | 3.1-2 |

**Trial-level Accuracy Analysis**

We modelled accuracy from the three-way interaction between group (patients, controls), hemifield of presentation (left, right), and trial type (valid, IO, IS), with mean-centered age as an additive effect and participant as a random effect. None of the interaction terms were significant. The only significant predictors of trial-level accuracy were age, as accuracy increased with older age (z = 2.26, p = 0.024); group, in which patients performed more poorly than controls overall (z = -3.12, p = 0.002); and trial type, in which accuracy to valid and invalid object trials was not statistically different (z = 1.11, p -0.27), but accuracy on invalid spatial trials was lower compared to both valid (z = -3.67, p = 0.0002) and invalid object trials (z = -3.56, p = 0.0004).

**Table S3**. Analysis of Deviance Table (Type II Wald chi-square tests) for parameters of accuracy model.

| **Model Term** | **Df** | **Χ^2^** | ***p*-value** | **BF** |
| --- | --- | --- | --- | --- |
| group x hemifield x trial type | 2 | 3.15 | 0.207 | 3966.27 |
| group x hemifield | 1 | 0.65 | 0.422 | 99.87 |
| trial type x hemifield | 2 | 1.78 | 0.410 | 7792.00 |
| group x trial type | 2 | 0.86 | 0.650 | 11984.92 |
| **group** | **1** | **10.72** | **0.001** | **0.17** |
| **trial type** | **2** | **24.69** | **<0.0001** | **1.05** |
| hemifield | 1 | 0.14 | 0.712 | 128.44 |
| **age** | **1** | **5.12** | **0.024** | **12.52** |

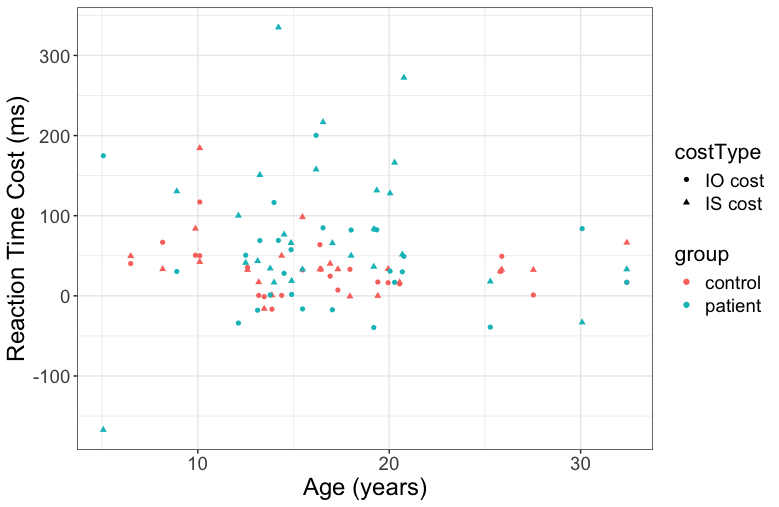

**Figure S1**. Relationship between age and average reaction time costs for IO (circle) and IO (triangle) trials for patients (teal) and controls (red).
